# Revisiting Macromolecular Hydration with HullRadSAS

**DOI:** 10.1101/2022.10.20.513022

**Authors:** Patrick J. Fleming, John J. Correia, Karen G. Fleming

## Abstract

Hydration of biological macromolecules is important for their stability and function. Historically, attempts have been made to describe the degree of macromolecular hydration using a single parameter over a narrow range of values. Here, we describe a method to calculate two types of hydration: surface shell water and entrained water. A consideration of these two types of hydration helps to explain the “hydration problem” in hydrodynamics. The combination of these two types of hydration allows accurate calculation of hydrodynamic volume and related macromolecular properties such as sedimentation and diffusion coefficients, intrinsic viscosities, and the concentration dependent non-ideality identified with sedimentation velocity experiments.

## Introduction

Biological macromolecules exist and function at least partly in an aqueous environment. Some of the solvent water is associated with each macromolecule and affects its stability and function; this water is considered macromolecular hydration. Many years ago, Kauzmann (Kauzmann, 1959) identified the hydrophobic effect as a primary force in the folding and stability of proteins and numerous studies have probed the nature of hydrophobic and polar hydration (Baldwin, 2014). Hydration water facilitates hydrogen bond switching and has been shown to be important in dynamical transitions in proteins (Tarek & Tobias, 2008; Dahanayake & Mitchell-Koch, 2018), in enzyme function (Rupley et al., 1983) and in the allosteric regulation of proteins (Colombo et al., 1992). The water of hydration must be displaced for binding of ligands and substrates to macromolecules and this process contributes favorable entropy for driving enzymatic function (Hwang et al., 2019). Water displacement is also a major driving force for macromolecular assembly of systems like microtubules (Lee & Timasheff, 1977; Vulevic & Correia, 1997).

Hydration also affects the hydrodynamic behavior of macromolecules; it increases the frictional drag for both translational and rotational diffusion. Diffusion and sedimentation of macromolecules is dependent not only on the size and shape of the macromolecule, but also on the amount of associated hydration water. Both diffusion and sedimentation coefficients are proportional to the translational hydrodynamic radius (*R_T_*) and it is convenient to use this characteristic parameter when comparing hydrodynamic properties as described by Garcia de la Torre (García de la Torre & Hernández Cifre, 2020). For a macromolecule with molecular mass *M*, the *R_T_* of an equivalent sphere with the same diffusion, or sedimentation, coefficient can be calculated from Eq. 1,

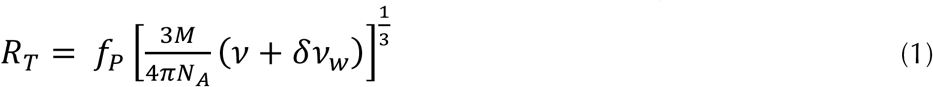

where *f*_p_ = a unitless frictional coefficient due to shape, *M* = molecular mass (g/mol), *ν* = macromolecular partial specific volume (ml/g), *δ* = amount of hydration water (g water/g macromolecule), *ν_w_* = hydration water partial specific volume (ml/g), and *N_A_* = Avogadro’s number. Assuming that the molecular mass and partial specific volumes are known, or can be estimated, it is necessary to know two variables to calculate the hydrodynamic radius: the frictional coefficient (*f*_p_) and the amount of hydration (*δ*). Currently, there is no way to solve unambiguously for both of them. This inability to separate the effects of hydration and shape has been called the “Hydration Problem” by Harding (Harding, 2001). If the macromolecular structure is known, a shape frictional coefficient may be estimated using Perrin’s equations (Cantor & Schimmel, 1980; Perrin, 1936), but as discussed below there is no universal value for the amount of hydration that can be used for all proteins or all nucleotides.

Many methods demonstrate that hydration water is associated with macromolecules. Over fifty years ago, Kuntz and colleagues used NMR to measure the hydration of polypeptides, each of which contained a specific amino acid type (poly-ALA, poly-ARG, etc.) (Kuntz, 1971). They assigned a hydration value to each of the naturally occurring amino acids and used these values, along with amino acid composition, to calculate the overall hydration of proteins. The four folded proteins studied were in the molecular weight range 15 - 65 kDa and had hydration of 0.31-0.45 g water/g protein. The calculated hydration, from amino acid composition, agreed remarkably well with experimentally determined hydration on folded proteins. This agreement with experimental values is surprising because it assumes that buried residues in folded proteins have the same hydration as homopolymers of the amino acid that are likely fully exposed to solvent. Nevertheless, numerous studies since that time on proteins of similar size have confirmed the same general range of hydration (Careri et al., 1980; Kuntz, 1971; Kuntz & Kauzmann, 1974; P. H. Yang & Rupley, 1979; Zhou, 1995).

Water molecules associated with proteins have been identified by X-ray crystallography. Most crystallographic waters in RNAse are on the surface in the first shell (Esposito et al., 2000) and represent a hydration of 0.34 g/g. It has also been demonstrated that different types of surface confinement influences hydration water properties (Persson et al., 2018). It is generally recognized that the above hydration, when distributed mostly in the first solvent shell represents a non-contiguous layer of water molecules. This first shell of hydration is idealized as a thin continuous surface layer for estimation of hydration volume (Cantor & Schimmel, 1980). In addition to first shell water, a second type of protein-associated water has been experimentally recognized and termed trapped water. This water is in small, buried cavities or narrow channels (Durchschlag & Zipper, 2003).

Despite general agreement that protein hydration is ~0.3-0.4 g/g, it is still not possible to use Eq. 1 to calculate accurate hydrodynamic properties such as diffusion coefficients or sedimentation coefficients for larger proteins using these general hydration values. As an alternative method to calculate hydrodynamic properties bead modeling of structural models does assume a general level of hydration, as reflected in the bead size (García de la Torre & Hernández Cifre, 2020) or by using the residue-based hydration of Kuntz as described above (Rocco & Byron, 2015). But these methods do not allow elucidation of the separate contributions of hydration and shape to the overall hydrodynamic behavior of the macromolecule. Although a common range of hydration levels may be sufficient to characterize small globular proteins, individual proteins with varied shape have additional associated water that affects hydrodynamic volume and it is this additional hydration water that is identified here.

Another method to calculate hydrodynamic properties from structural models is HullRad. HullRad is a computer program that uses a convex hull around a macromolecule to determine the molecular hydrodynamic volume (Fleming & Fleming, 2018). We have implemented a new method into the HullRad algorithm that allows calculation of the specific hydration of macromolecules. The method is implemented in a computer program called HullRadSAS. Analysis of the hydration associated with a variety of proteins and DNA demonstrates that water *entrained* by the macromolecule during diffusion, but not necessarily in contact with the macromolecular surface, contributes to the hydrodynamic volume of the macromolecule. Large proteins have significantly more entrained water than small proteins. Inclusion of entrained water in the total hydration (*δ*) allows accurate calculation of translational and rotational hydrodynamic properties.

It should be noted that Creeth and Knight used the term “solvent entrainment” to describe protein hydration (Creeth & Knight, 1965). As they defined the term, it did not connote any distinction between “tightly bound” and “loosely bound” solvent. Here, we are specifically using *entrainment* to indicate the water within the convex hull but not on the surface of the macromolecule, and *shell* to indicate the hydration sites on the surface of a macromolecule always occupied by a water molecule. Our use of the terms *shell* and *entrainment* is consistent with the description by Harding that hydration “...represents the amount of solvent ‘associated’ with the macromolecule and includes ‘chemically bound’ via hydrogen bonds and ‘physically entrained’ solvent” (Harding, 1997).

## Methods

The design of HullRadSAS differs from the original HullRad (Fleming & Fleming, 2018) as illustrated in Fig. 1. For both versions a coarse-grained model of the macromolecule containing backbone atoms and a single pseudo-atom side chain for each residue is built (Fig. 1, A and B, Left panel). This coarse-graining effectively averages residue side chain rotamers. Similar coarse-graining is done for polynucleotides and oligosaccharides. In the original HullRad an initial convex hull is constructed on the atom sphere centers of the coarse-grained model (Fig. 1, A, Middle panel). The planes of this initial hull are expanded 2.8 Å along each plane normal to account for hydration (Fig. 1, A, Right panel).

**FIGURE 1.**
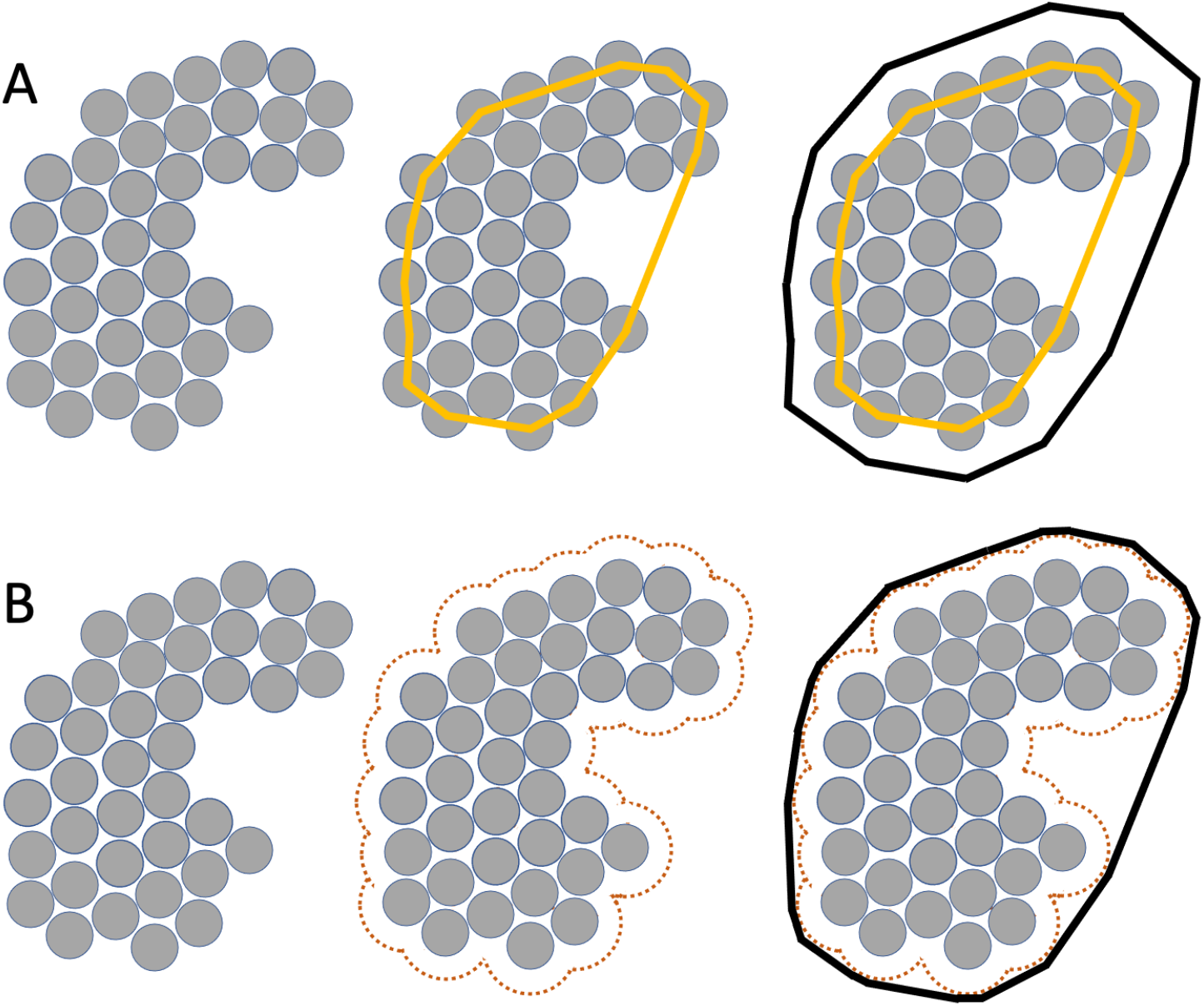
Design of HullRad and HullRadSAS. The figure is a two-dimensional representation of the three-dimensional objects. A, Original HullRad; Left, coarse-grained model of the macromolecule (gray spheres). Middle, a convex hull (orange lines) constructed on the atom sphere centers of the coarse-grained model. Right, Initial convex hull is expanded to account for hydration (black lines). B, HullRadSAS; Left, coarsegrained model of the macromolecule (gray spheres). Middle, a solvent accessible surface (orange dots) constructed using the sphere centers of the coarse-grained model. Right, a convex hull (black lines) constructed to enclose the points of the accessible surface. The radius of a sphere equivalent to the volume of the black convex hulls is proportional to the hydrodynamic volume of the macromolecule.

In HullRadSAS a solvent accessible surface (SAS) is constructed using the sphere centers of the coarse-grained model (Fig. 1, B, Middle panel). The points of the accessible surface are then used to construct a final convex hull (Fig 1, B, Right panel). The final convex hull constructed this way is analogous to the expanded convex hull described for the original HullRad. An ellipsoid of revolution is built with a major axis equal to the maximum dimension and the same volume of the convex hull; the major and minor axes of this ellipsoid are then used to calculate a frictional shape factor. The radius of a sphere equivalent to the volume of the convex hull, multiplied by the shape factor, equals the translational hydrodynamic radius of the macromolecule.

HullRadSAS is implemented in Python using the SASA module from Biopython (Cock et al., 2009). The code is freely available from the HullRad website and GitHub.

## Results and Discussion

### Hydration and Translational Hydrodynamics

To validate HullRadSAS, we compared the performance of HullRadSAS to the original HullRad. Both versions of the program calculate essentially identical results for the translational hydrodynamic radii (*R*_T_) of 32 proteins from the data set used to calibrate HullRad (Fleming & Fleming, 2018) (Supplemental Fig. S1 and Supplemental Table S1).

In HullRadSAS an SAS of the macromolecule is used as the object for construction of a convex hull. The optimal value of 0.85 Å for SAS probe radius was determined empirically (Supplemental Fig. S2). In the original HullRad the planes of the convex initial hull were expanded 2.8 Å. This expansion of the initial hull is to account for hydration and the optimal expansion of 2.8 Å was empirically determined (see Supplemental Figure 3 in Fleming and Fleming (Fleming & Fleming, 2018)). In HullRadSAS the convex hull is a priori expanded because it is constructed on the surface points of the SAS. Here the distance between the convex hull and the atom centers of the coarse-grained model equals 0.85 Å plus the radius of the closest atom. Atoms in the coarse-grained model include both back-bone atoms and the side chain single pseudo-atoms, all with atomic radius arbitrarily defined as 2.0 Å for calculation of the SAS. Therefore, in HullRadSAS, the distance between the hull planes and closest atom center is 0.85 + 2.0 = 2.85 Å. We note that the hydrodynamic prediction program BEST also uses an SAS in the algorithm (Aragon & Hahn, 2006). In this case the SAS is used as a basis for computing boundary elements.

Hydrodynamic volume is defined as the sum of the time-average of the molecular volume and the volume of the solvent molecules associated with it (Cammack et al., 2006). Fig. 2 illustrates two types of water that contribute to the hydrodynamic volume of a macromolecule: Surface shell water and entrained water. In the procedure described here the first shell of hydration (defined by the SAS) is modeled by a continuous layer. The amount of water in the shell volume is calculated using a density of water 10% greater than that of bulk water (Halle, 2004). The amount of water in the entrained volume is calculated using the density of bulk water.

**FIGURE 2.**
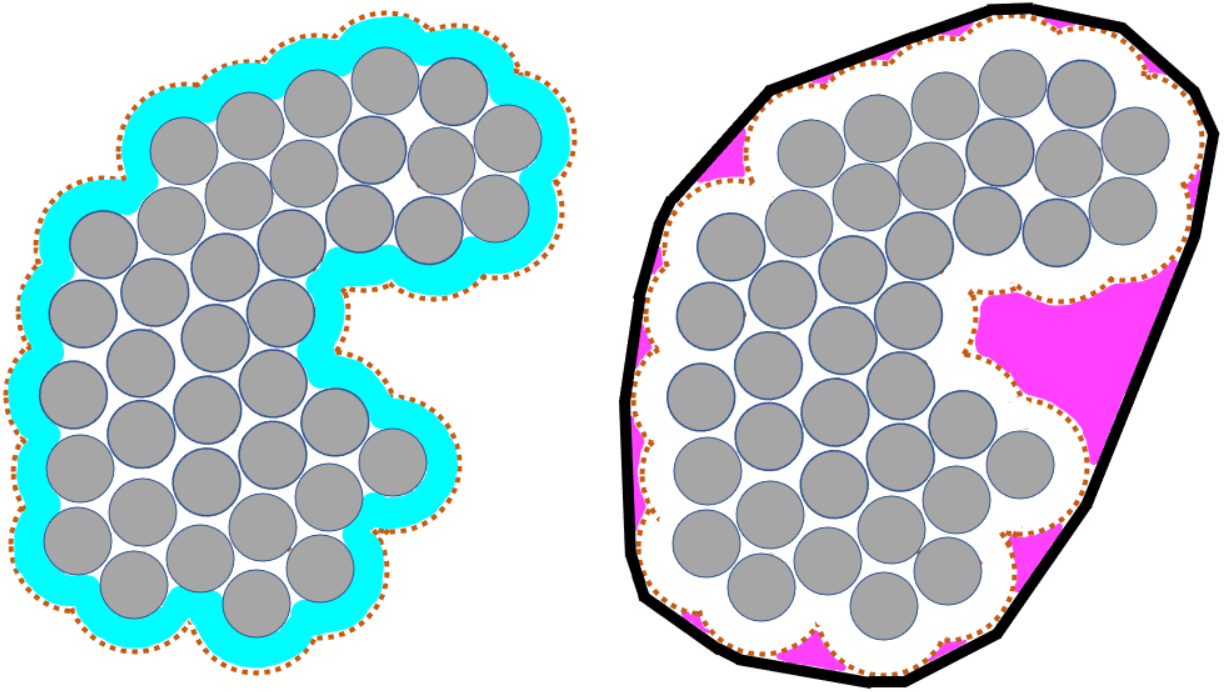
HullRadSAS provides the means to calculate two types of hydration. The figure is a two-dimensional representation of the three-dimensional objects. Left, the volume (cyan) between the SAS (orange dots) and surface of the macromolecular atoms (gray spheres) represents the first hydration shell. Right, the volume (magenta) between the SAS (orange dots) and convex hull (thick black lines) represents water entrained in crevices, grooves, and pockets of the molecule and part of the hydrodynamic volume.

Only the first shell water (plus a small amount of second shell water) is observed with methods such as NMR (Esposito et al., 2000). As calculated by HullRadSAS the amount of shell water is between 0.23-0.35 g water/g protein for a set of proteins ranging in molecular weight from 6 to 828 kDa (Fig. 3). The amount of shell water is proportional to protein size for small proteins but is consistently between 0.23 and 0.28 g/g for proteins larger than ~200 kDa (Fig. 3). As discussed below, larger proteins have a more rugged surface topology or extended non-spherical shape with proportionately more surface area/volume and these factors account for the fact that first shell hydration does not follow the expected decrease in surface area to volume ratio of a sphere for these larger proteins.

**FIGURE 3.**
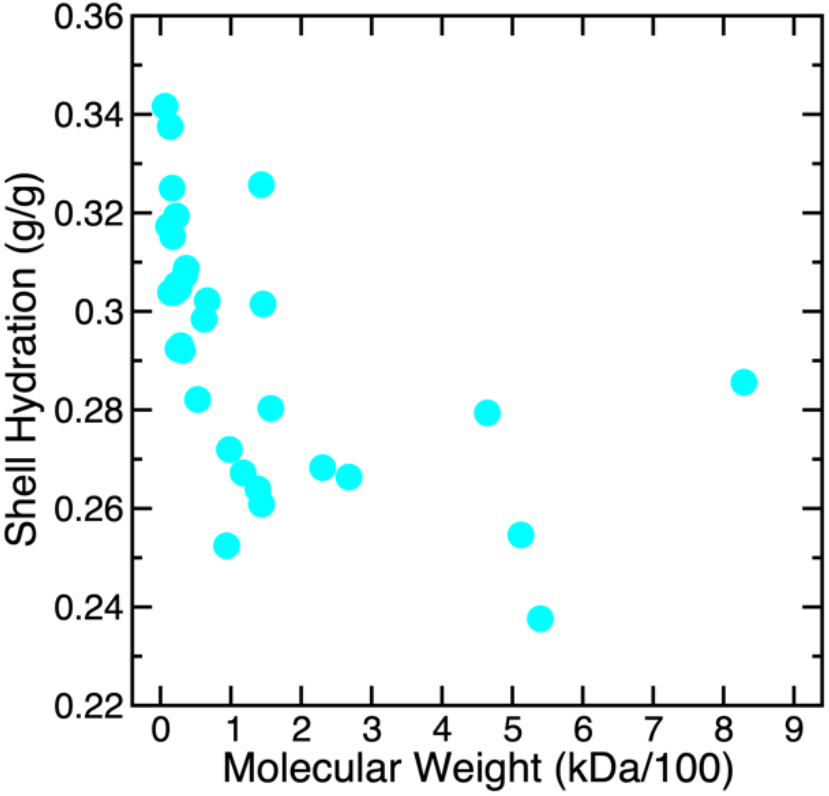
HullRadSAS calculated first shell hydration water depends on protein size. The amount of shell water calculated by HullRadSAS is plotted versus protein molecular weight as cyan circles.

For small globular proteins the first shell water makes up most of the macromolecular hydration, but larger proteins have significantly more entrained water (Fig. 4). Proteins larger than ~200 kDa have 0.5 g/g or more of entrained water. Two proteins in the data set, IgG and GroEL, include extraordinary amounts of entrained water and the reason for this is illustrated in Fig. 5. IgG has large spaces between the domains that are encapsulated within the convex hull, and GroEl has a large cavity that is completely within the protein itself. It is important to re-iterate that the volume of the convex hull is proportional to the hydrodynamic volume of these proteins, so the entrained water is effectively part of the protein during diffusion and sedimentation.

**FIGURE 4.**
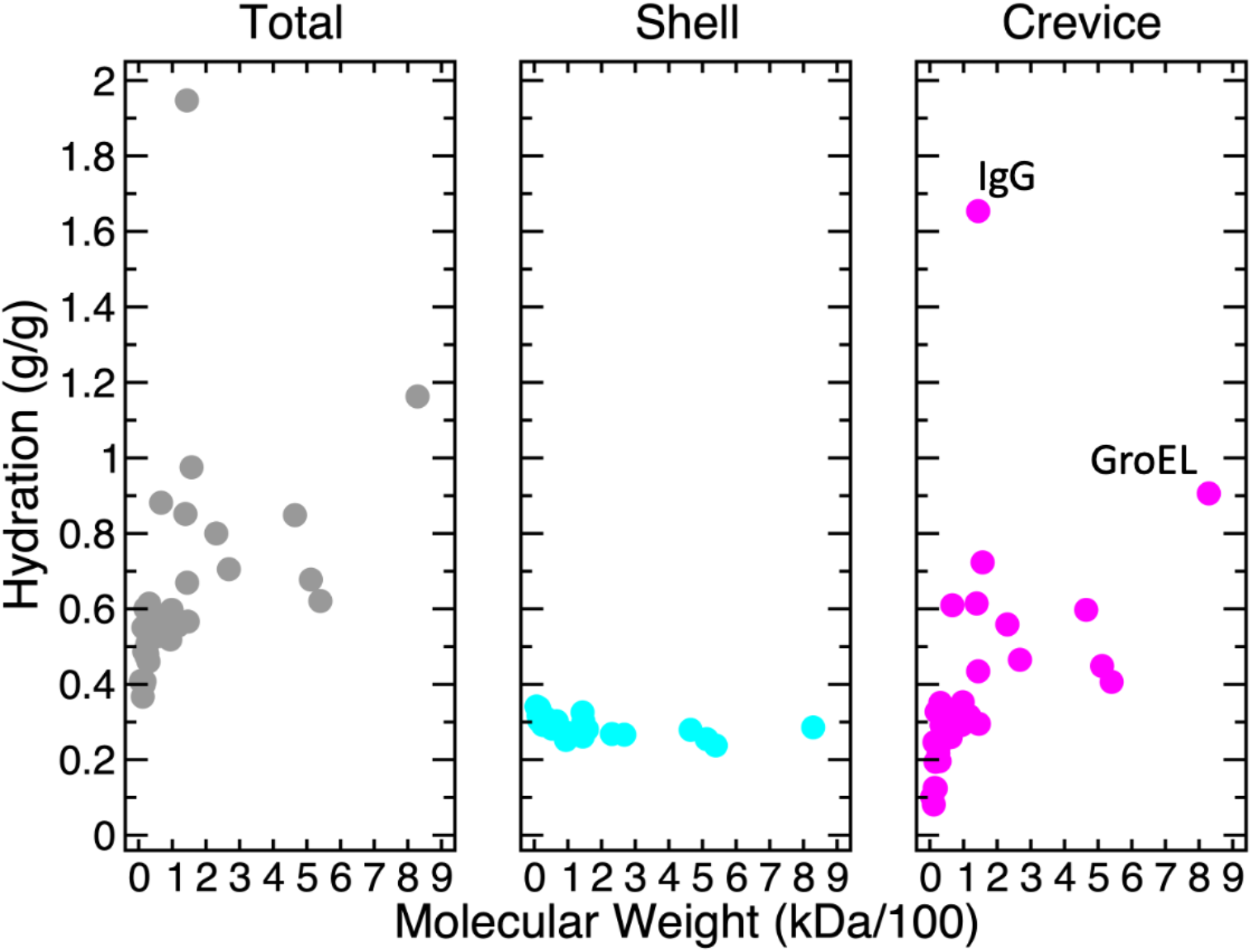
HullRadSAS calculates both shell and entrained hydration water. Left panel, total hydration water for the data set of proteins in Supplemental Table S1 is plotted versus protein molecular weight as grey circles; middle panel, first shell hydration water amount is plotted as cyan circles (the data is the same as in Fig. 3); right panel, the entrained water is plotted as magenta circles. Two proteins with large amounts of entrained water are labeled (IgG and GroEL).

**FIGURE 5.**
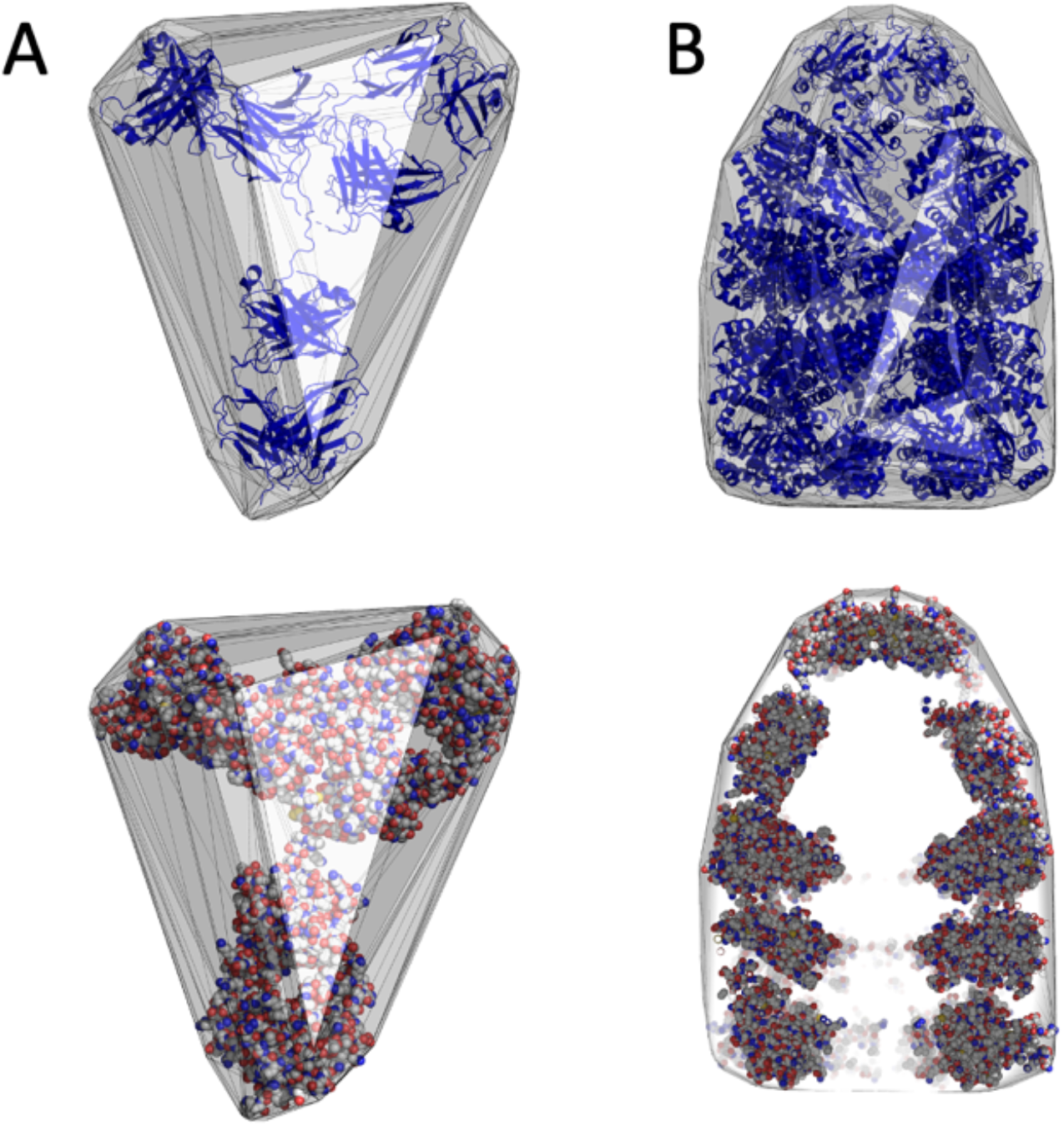
Some proteins contain large crevices or cavities within the hydrodynamic volume. A, Human IgG (PDBcode: 1HZH) and B, *E. coli* GroEL (PDBcode: 2CGT) are shown as ribbon drawings (top) and atomic sphere models (bottom) together with their respective convex hulls. GroEL within the convex hull in the bottom image is shown as a slice to visualize the internal cavity. The convex hulls were constructed on the respective SAS points for each protein. PyMOL (DeLano, 2015) was used to create the images.

The convex hull constructed by the HullRad algorithm accurately describes the hydrodynamic volume of greatly expanded structures found in the structural ensemble of an intrinsically disordered protein (IDP) (Fleming & Fleming, 2018). Inevitably, large amounts of entrained water are encapsulated by the convex hull of an expanded protein. Differential residue hydration has been found to explain the sequence dependence of IDP expansion/collapse (Wuttke et al., 2014), but these sequence specific effects would be expected to influence first shell hydration and not entrained hydration. In addition, surface shell water may be subtly different for unfolded proteins compared to the native folded state (Sengupta et al., 2008). However, the degree of expansion, partially driven by sequence specific hydration, is hydrodynamically modeled by HullRad.

The inclusion of hydration water beyond the first shell in the hydrodynamic volume is demonstrated in more detail by consideration of the data in Table 1 and Fig. 6. The first four proteins listed in Table 1 are relatively small proteins with shell hydration of 0.30-0.34 g/g and entrained hydration of 0.12-0.25 g/g. Fig. 6 (top panel) shows that these four proteins are well described as ellipsoidal in shape and without major crevices or grooves. In contrast, the last four proteins have similar, or slightly less, shell hydration but much larger entrained hydration of 0.41 −1.65 g/g. Fig. 6 (bottom panel) shows that these latter four proteins have tortuous surfaces with large crevices or grooves. It is only with the inclusion of the entrained water in the hydrodynamic volume that accurate hydrodynamic radii are calculated (Table 1, last column).

**FIGURE 6.**
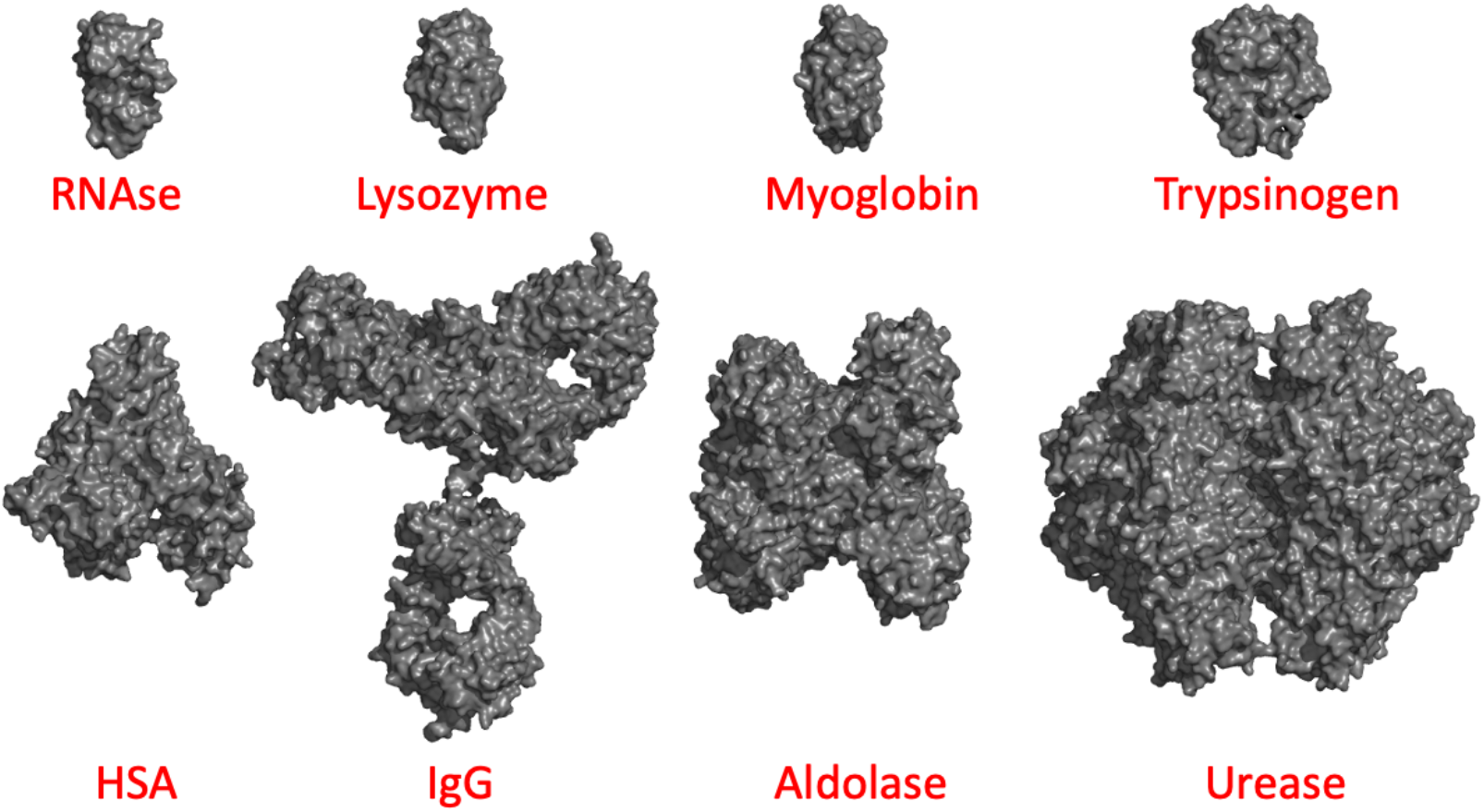
Many large proteins are non-spherical in shape with tortuous surfaces. The eight proteins listed in TABLE 1 are shown as surface models, all images are to the same scale. PyMOL (DeLano, 2015) was used to create the images.

**Table 1.** Hydration and Translational Hydrodynamic Radii for Folded Proteins.

| Protein (PDB Code) | MW (Da) | $R_{T,Exp}^a$ (Å) | Shell $\delta^b$ (g/g) | Entrained $\delta^b$ (g/g) | $R_{T,HRSAS}^c$ (Å) | $R_{T,HRSAS} - R_{T,Exp}$ (%Error) |
| --- | --- | --- | --- | --- | --- | --- |
| RNase A (8RAT) | 13692 | 18.9 | 0.34 | 0.25 | 19.0 | 0.58 |
| Lysozyme (1AKI) | 14315 | 18.6 | 0.30 | 0.13 | 18.6 | -0.16 |
| Apo-Mb (1MGN) | 17359 | 20.8 | 0.32 | 0.12 | 20.0 | -3.71 |
| Trypsinogen (1TGN) | 22620 | 22.2 | 0.32 | 0.20 | 22.2 | 0.21 |
| HSA (1AO6) | 66482 | 34.0 | 0.30 | 0.61 | 35.0 | 3.06 |
| Human IgG (1HZH) | 143337 | 55.1 | 0.33 | 1.65 | 54.6 | -0.93 |
| Aldolase (1ADO) | 156776 | 47.6 | 0.28 | 0.72 | 47.6 | -0.16 |
| Urease (3LA4) | 539700 | 65.8 | 0.24 | 0.41 | 66.4 | 0.91 |

<sup>a</sup>Experimental $R_T$ values from Supplemental Table S3.
<sup>b</sup>Shell and entrained hydration water calculated with HullRadSAS.
<sup>c</sup> $R_T$ calculated with HullRadSAS.

In contrast to the consensus described in the Introduction that proteins have a common and limited range of hydration, Squire and Himmel provided a detailed analysis for a set of proteins with known structures and concluded that “individual proteins demonstrate wide variations in their hydration levels” (Squire & Himmel, 1979). They went on to say that these variations reflect “considerable individual character” with respect to transport properties. Our data shows that the tortuous surfaces with varying amounts of entrained water of proteins illustrated in the bottom panel of Fig. 6 contribute to the individual character of proteins found by transport methods.

### Hydration water dynamics

The fact that entrained water is part of the hydrodynamic volume does not mean that the same water molecules are permanently associated with the protein. Residence times for water molecules in the first shell of hydration are on the order of tens of ps (Halle & Davidovic, 2003; Wüthrich et al., 1996). But entrained water in crevices is expected to have the mobility of bulk water with diffusion coefficients at least 10-100 times larger (Rupley & Careri, 1991). With a “residence time” in a crevice of <1 ps, entrained water will diffuse in and out of a crevice many times while a protein diffuses the distance of its diameter. However, entrained water statistically will be part of the macromolecule as the protein diffuses. As stated by Halle and Davidovic, “...large-scale shape irregularities, such as [a] binding cleft …, make the [diffusing] protein displace a larger amount of solvent than would a compact protein of the same volume” (Halle & Davidovic, 2003).

The geometrical definition of entrained water illustrated in Fig. 2 is that water enclosed by the convex hull minus shell water. Therefore, true “trapped” water, that water with long residence times in internal cavities would be included in the geometrical definition of entrained water. Although the amount of water in internal cavities usually is small relative to the total hydration described here (Williams et al., 1994), it would be included in the entrained category and, of course, be part of the diffusing macromolecule.

### Non-ideality in sedimentation velocity analysis

Creeth and Knight clearly envisioned that the amount of entrained water could vary with the protein (Creeth & Knight, 1965). They argued that large effective hydrodynamic volumes from extensive solvent entrainment provided an interpretation of concentrationdependent non-ideality revealed in sedimentation velocity experiments.

Sedimentation coefficients are usually obtained at finite concentrations of solute and corrected to zero concentration. This is necessary because the measured sedimentation coefficient (*s*) has been found to be concentration dependent according to the following generally accepted relationship,

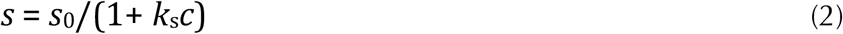

where *S*_0_ is the sedimentation coefficient at infinite dilution, *c* is the solute concentration, and *k*_s_ is an empirically determined constant known as the hydrodynamic non-ideality constant. Rowe derived the following relationship between *k*_s_ and hydrodynamic properties (Eq. 11 in Rowe (Rowe, 1977)),

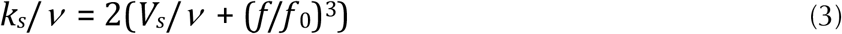

where *n* is the partial specific volume of the macromolecule (ml/g), (*f/f*_0_)^3^ is a unitless effective frictional ratio comprised of two factors due to swelling and asymmetry (Eq. 22 in Rowe (Rowe, 1977)), and *V_s_* is the specific volume associated with the unit mass of hydrated macromolecules (obtainable from the convex hull volume calculated by HullRad and HullRadSAS).

Concentration dependence in sedimentation velocity experiments may be due to macromolecular hydration, shape asymmetry, or interactions such as self-association or association with other components in solution. Global modeling of these effects in high concentration solutions is now possible with analytical ultracentrifuge data analysis software such as SEDANAL (Stafford, 2016; Stafford & Sherwood, 2004). The range of *k*_s_ found experimentally is from ~2 to 20 ml/g for globular proteins (Wright et al., 2018; Creeth & Knight, 1965). Wright et al. (Wright et al., 2018) studied the non-ideality of a monoclonal antibody and human IgG with sedimentation velocity experiments and determined *k_s_* values of ~3.0 ml/g. When they accounted for the effect of self-association in the fitting model with SEDANAL the *k_s_* due only to hydration and shape was increased to ~11 ml/g. When Eq. 3 is implemented in HullRadSAS the calculated *k_s_* for human IgG is 11.1 ml/g. This striking agreement lends support to the results reported in Wright et al. (Wright et al., 2018) and to the conclusion of Yang et al. (D. Yang et al., 2018) that experimental values of *k_s_* that deviate from calculated values are suggestive of weak association that masks the magnitude of *k_s_*.

A comparison of calculated and experimental *k_s_* values in concentrated and/or complicated solutions would help identify interacting systems. Such information would be useful to understand the influence of serum or crowded cellular compartments on the diffusion properties of macromolecules.

### Rotational Hydrodynamics and Intrinsic Viscosity

The basis of the HullRad algorithm is that an expanded convex hull represents the hydrated hydrodynamic volume of a macromolecule. As described above the *translational* hydrodynamic radius of a macromolecule may be calculated by multiplying the radius of a sphere equivalent to the volume of the expanded convex hull by a shape factor. In this case, the shape factor is derived from Perrin’s equations.

As described in the Supplemental Information for Fleming and Fleming (Fleming & Fleming, 2018), the hydration layer has different effects on *rotational* and *translational* diffusion, and rotational diffusion is affected by asymmetry differently than translational diffusion. In the original HullRad these differences for rotational diffusion are accounted for by empirical adjustment of both the hydration layer thickness and Perrin-derived shape factor. HullRadSAS uses the same approach and also calculates accurate rotational hydrodynamic properties. The hydration and rotational hydrodynamic radii for four small and four large proteins are listed in Table 2. All these proteins have shell hydration of 0.33 - 0.35 g/g. The four smallest proteins have small entrained hydration of 0.06 - 0.1 g/g, and the larger proteins have entrained hydration of 0.16 - 0.51 g/g. Again, inclusion of the entrained hydration volumes is necessary to obtain calculated hydrodynamic radii that agree with experimental values (Table 2, last column).

**Table 2.** Hydration and Rotational Hydrodynamic Radii for Folded Proteins.

| Protein (PDB Code) | MW (Da) | $R_{R,Exp}^a$ (Å) | Shell $\delta^b$ (g/g) | Entrained $\delta^b$ (g/g) | $R_{R,HRSAS}^c$ (Å) | $R_{R,HRSAS} - R_{R,Exp}$ (%Error) |
| --- | --- | --- | --- | --- | --- | --- |
| BPTI (5PTI) | 6518 | 15.3 | 0.34 | 0.10 | 15.9 | 3.67 |
| Calbindin (1IG5) | 8502 | 17.0 | 0.35 | 0.10 | 16.9 | -0.53 |
| Ubiquitin (1UBQ) | 8566 | 16.8 | 0.35 | 0.10 | 17.4 | 3.27 |
| Plastocyanin (1PCS) | 10322 | 18.0 | 0.33 | 0.06 | 17.8 | -0.78 |
| RNase A (1AQP) | 13692 | 20.0 | 0.33 | 0.25 | 20.6 | 3.10 |
| Spo0F (2FSP) | 14230 | 20.1 | 0.33 | 0.16 | 20.5 | 2.04 |
| β-Lactglob (2AKQ) | 17963 | 22.6 | 0.33 | 0.23 | 22.5 | -1.49 |
| Apo-AK (4AKE) | 23589 | 26.1 | 0.33 | 0.51 | 26.7 | 3.25 |

<sup>a</sup>Experimental $R_R$ values from Table 2 in Fleming and Fleming (Fleming & Fleming, 2018).
<sup>b</sup>Shell and entrained hydration water calculated with HullRadSAS.
<sup>c</sup> $R_R$ calculated with HullRadSAS.

Intrinsic viscosity may also be accounted for in terms of hydrodynamic volume and shape asymmetry (Rowe, 1977). The intrinsic viscosity ([η]) of a sphere is related to the hydrodynamic volume (*V*_h_) as described by Einstein,

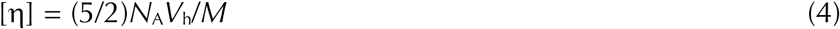

where *N_A_* and *M* are defined above for Eq. 1. For asymmetric macromolecules [η] is exquisitely sensitive to shape especially for large axial ratios and this sensitivity makes intrinsic velocity a useful property for describing elongated and rod-like macromolecules. To account for shape effects the factor of (5/2) in Eq. 4 is sometimes replaced by the Simha factor (Cantor & Schimmel, 1980; Rowe, 1977) for calculating intrinsic viscosity. We have found that a function derived from the gyration tensor alone (García de la Torre et al., 2000) works well as a shape factor to calculate intrinsic viscosity. Supplemental Table S2 and Supplemental Fig. S3 illustrate the excellent agreement between the intrinsic viscosities calculated by HullRadSAS and experimental values for a set of proteins and DNA duplexes.

### Hydration and Nucleic Acids

The data in Table 3 indicate that duplex DNA has slightly more first shell hydration (0.35-0.38 g/g) than the proteins listed in Tables 1 and 2. The entrained water in B-DNA represents significantly more hydration with values ranging from 0.47-0.70 g/g that are proportional to the length of duplex (Table 3). The slight length dependence on the amount of entrained water is due to end effects for these relatively short duplexes, and the relative proportion of entrained water in the grooves would be expected to level off for very long duplexes where end effects become insignificant. Z-DNA has similar first shell hydration (0.37 g/g), but the entrained hydration is less than found for B-DNA of the same length. G-quadruplex DNA also has a similar amount of shell water (0.36 g/g) and an intermediate level of entrained water. It should be noted that the duplex DNA examples in Table 3 would be too short to exhibit flexible bending and therefore a single structure is representative of the solution conformation (Kovacic & van Holde, 1977). Fig. 7 illustrates that both the minor and major grooves of B-DNA (Panel A) contribute significant volume encapsulated by the convex hull whereas the shallow grooves of Z-DNA (Panel B) contribute much less encapsulated volume. G-quadruplex-DNA (Panel C) has fewer large crevices compared to B-DNA and this is responsible for the intermediate level of entrained water.

**FIGURE 7.**
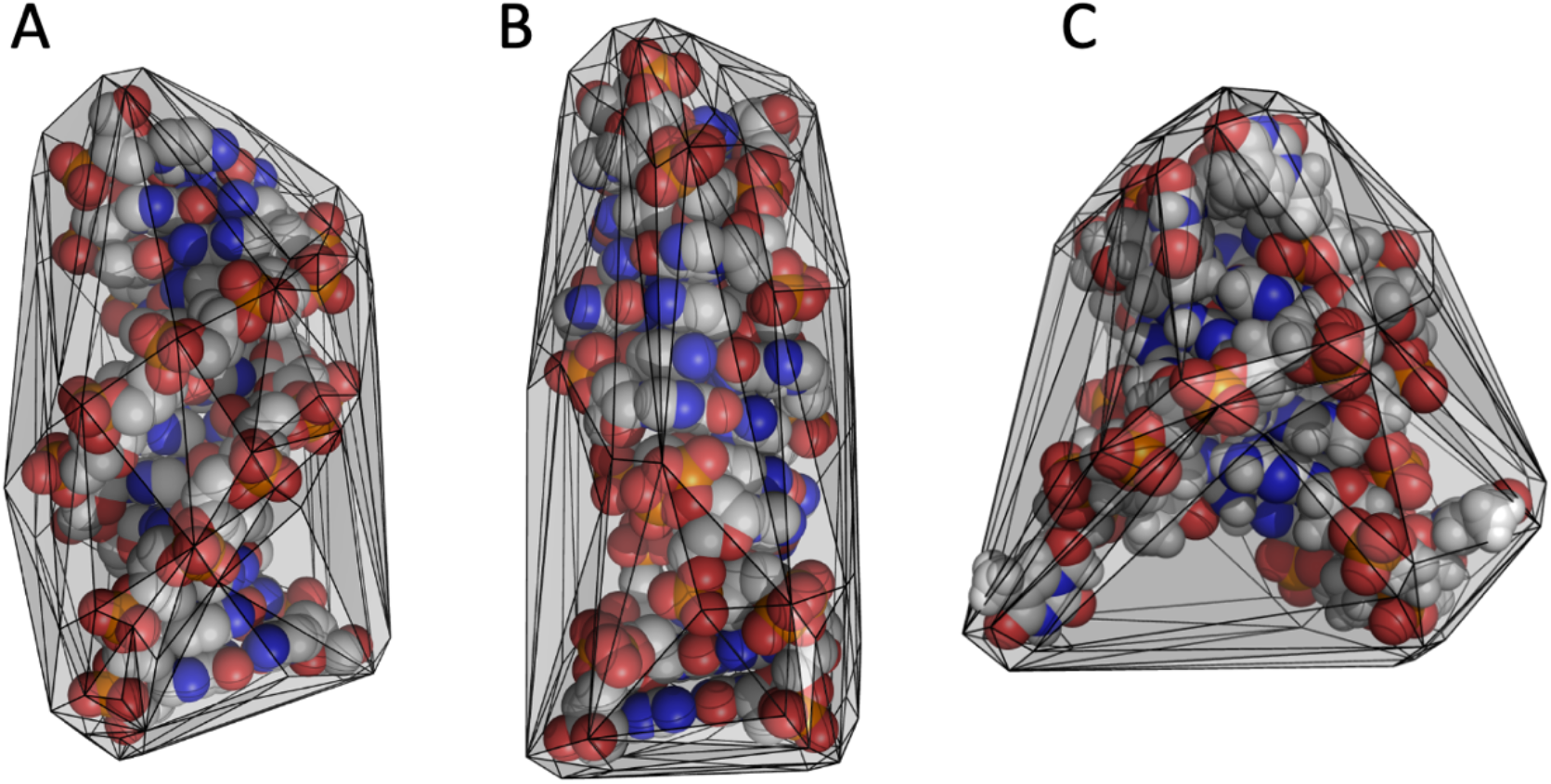
Different DNA structures have different hydration volumes. A, 12 bp B-DNA duplex; B, Z-DNA duplex; and C, Q-quadruplex-DNA are shown as atomic spheres. The respective convex hulls constructed by HullRadSAS are shown with gray planes and edges as black lines. PyMOL (DeLano, 2015) was used to create the images. See Table 3 for hydration details.

**Table 3.** Hydration and Translational Hydrodynamic Radii for DNA.

| Protein (PDB Code) | MW (Da) | $R_{T,Exp}$ (Å) | Shell $\delta^b$ (g/g) | Entrained $\delta^b$ (g/g) | $R_{T,HRSAS}^c$ (Å) | $R_{T,HRSAS} - R_{T,Exp}$ (%Error) |
| --- | --- | --- | --- | --- | --- | --- |
| B-DNA 8mer | 4832 | 14.1<sup>a</sup> | 0.38 | 0.47 | 14.0 | -0.01 |
| B-DNA 12mer | 7313 | 16.6<sup>a</sup> | 0.37 | 0.55 | 16.5 | -0.01 |
| B-DNA 20mer | 12270 | 19.7<sup>a</sup> | 0.35 | 0.66 | 20.4 | 0.04 |
| B-DNA 24mer | 15216 | 22.5<sup>a</sup> | 0.36 | 0.70 | 22.4 | -0.01 |
| Z-DNA 12mer (4OCB) | 7319 | N.A. | 0.37 | 0.32 | 15.7 | N.A. |
| G-quadruplex (143D<sup>d</sup>) | 6988 | 15.8<sup>e</sup> | 0.36 | 0.36 | 15.3 | -2.84 |

<sup>a</sup>Experimental $R_T$ values from Table 3 in Fleming and Fleming (Fleming & Fleming, 2018).
<sup>b</sup>Shell and entrained hydration water calculated with HullRadSAS.
<sup>c</sup> $R_T$ calculated with HullRadSAS.
<sup>d</sup>NMR model 4, with lowest RMSD from ensemble average was used.
<sup>e</sup>Experimental $R_T$ value is average calculated from data in Li et al. (Li, 2005) and Hellman et al. (Hellman et al., 2010) using a partial specific volume of 0.525 as determined by Hellman et al. (Hellman et al., 2010)

An infrared study of protein-free calf thymus DNA indicated on the order of 20 molecules of water per nucleotide associated with B-DNA and with only 5-6 tightly bound (Falk et al., 1963). This total number of water molecules would represent slightly greater than 1 g/g of hydration and agrees with both gravimetric (Falk et al., 1962) and neutron quasielastic scattering methods (Bastos et al., 2004). Consistent with these infrared results, neutron scattering identified two types of water: ~30% strongly attached to the B-DNA surface with the remainder having a limited diffusive motion (Bastos et al., 2004). In contrast, only a total of 9 water molecules per nucleotide are found to fully hydrate Z-DNA (Umehara et al., 1990). The hydration of both forms of DNA calculated by HullRadSAS is consistent with these accumulated experimental results that describe the two types, and relative amounts, of hydration.

## Summary

A method using the solvent accessible surface to construct a convex hull around a macromolecule for calculating the hydrodynamic volume is described. The method allows differentiation of two types of macromolecular hydration: Surface shell and entrained water. Surface hydration calculated by this method agrees with historical values of ~0.35 g/g which largely measured only the first shell hydration. Larger proteins and DNA have significantly more entrained water (0.5 - 1.6 g/g) that contributes to hydrodynamic volume. The hydrodynamic volumes calculated by HullRad and HullRadSAS allow accurate calculation of several hydrodynamic properties including: translational and rotational hydrodynamic radii, intrinsic viscosities, and the concentration dependence of the sedimentation constant due to hydration. While only HullRadSAS provides hydration values, both versions calculate the same macromolecular hydrodynamic volume and therefore the same hydrodynamic properties. HullRadSAS is slower than the original HullRad because of the need to calculate the solvent accessible surface.

## Supplementary Information

The online version contains supplementary material.

## Authors contributions

P.J.F. designed the work and wrote the code; J.J.C. contributed the ideas on non-ideality; P.J.F., J.J.C. and K.G.F. wrote the manuscript. All authors read and approved the final manuscript.

## Availability of data and materials

All data are included in the text or supplementary information.

## Code availability

Code for both HullRadSAS and HullRad is available at the HullRad website (hullrad.wordpress.com, hullrad.jhu.edu) and GitHub (github.com/fleming12/hullrad.github.io).

## Acknowledgements

We would like to thank Dr. Barbara Amann for suggesting a comparison of B- and Z-DNA hydration and all members of the Fleming lab for helpful discussions.

## References

Aragon, S., & Hahn, D. K. (2006). Precise Boundary Element Computation of Protein Transport Properties: Diffusion Tensors, Specific Volume, and Hydration. Biophysical Journal, 91(5), 1591–1603. https://doi.org/10.1529/biophysj.105.078188

Baldwin, R. L. (2014). Dynamic hydration shell restores Kauzmann’s 1959 explanation of how the hydrophobic factor drives protein folding. Proceedings of the National Academy of Sciences of the United States of America, 111(36), 13052–13056. https://doi.org/10.1073/pnas.1414556111

Bastos, M., Castro, V., Mrevlishvili, G., & Teixeira, J. (2004). Hydration of ds-DNA and ss-DNA by Neutron Quasielastic Scattering. Biophysical Journal, 86(6), 3822–3827. https://doi.org/10.1529/biophysj.104.039586

Cammack, R., Atwood, T., Campbell, P., Parish, H., Smith, A., Vella, F., & Stirling, J. (Eds.). (2006). Oxford Dictionary of Biochemistry and Molecular Biology. Oxford University Press. https://doi.org/10.1093/acref/9780198529170.001.0001

Cantor, C. R., & Schimmel, P. R. (1980). Biophysical Chemistry, Part 2: Techniques for the Study of Biological Structure and Function. W.H. Freeman and Company.

Careri, G., Grattont, E., Yangt, P., & RupleyH, J. A. (1980). Correlation of IR spectroscopic, heat capacity, diamagnetic susceptibility and enzymatic measurements on Iysozyme powder. Proc. Nat. Acad. Sci. U.S.A, 284(9), 7955–7958.

Cock, P. J. A., Antao, T., Chang, J. T., Chapman, B. A., Cox, C. J., Dalke, A., Friedberg, I., Hamelryck, T., Kauff, F., Wilczynski, B., & de Hoon, M. J. L. (2009). Biopython: Freely available Python tools for computational molecular biology and bioinformatics. Bioinformatics, 25(11), 1422–1423. https://doi.org/10.1093/bioinformatics/btp163

Colombo, M. F., Rau, D. C., & Parsegian, V. A. (1992). Protein Solvation in Allosteric Regulation: A Water Effect on Hemoglobin. Science, 256(5057), 655–659. https://doi.org/10.1126/science.1585178

Creeth, J. M., & Knight, C. G. (1965). On the estimation of the shape of macromolecules from sedimentation and viscosity measurements. Biochimica et Biophysica Acta (BBA) - Biophysics Including Photosynthesis, 102(2), 549–558. https://doi.org/10.1016/0926-6585(65)90145-7

Dahanayake, J. N., & Mitchell-Koch, K. R. (2018). Entropy connects water structure and dynamics in protein hydration layer. Physical Chemistry Chemical Physics, 20(21), 14765–14777. https://doi.org/10.1039/C8CP01674G

DeLano, W. L. (2015). The PyMOL Molecular Graphics System (2.5.0). Schrodinger, LLC.

Durchschlag, H., & Zipper, P. (2003). Modeling the hydration of proteins: Prediction of structural and hydrodynamic parameters from X-ray diffraction and scattering data. European Biophysics Journal, 32(5), 487–502. https://doi.org/10.1007/s00249-003-0293-z

Esposito, L., Vitagliano, L., Sica, F., Sorrentino, G., Zagari, A., & Mazzarella, L. (2000). The ultrahigh resolution crystal structure of ribonuclease A containing an isoaspartyl residue: Hydration and stereochemical analysis. Journal of Molecular Biology, 297(3), 713–732. https://doi.org/10.1006/jmbi.2000.3597

Falk, Michael., Hartman, K. A., & Lord, R. C. (1962). Hydration of Deoxyribonucleic Acid. I. a Gravimetric Study. Journal of the American Chemical Society, 84(20), 3843–3846. https://doi.org/10.1021/ja00879a012

Falk, Michael., Hartman, K. A., & Lord, R. C. (1963). Hydration of Deoxyribonucleic Acid. II. An Infrared Study. Journal of the American Chemical Society, 85(4), 387–391. https://doi.org/10.1021/ja00887a004

Fleming, P. J., & Fleming, K. G. (2018). HullRad: Fast Calculations of Folded and Disordered Protein and Nucleic Acid Hydrodynamic Properties. Biophysical Journal, 114(4), 856–869. https://doi.org/10.1016/j.bpj.2018.01.002

García de la Torre, J., & Hernández Cifre, J. G. (2020). Hydrodynamic Properties of Biomacromolecules and Macromolecular Complexes: Concepts and Methods. A Tutorial Mini-review. Journal of Molecular Biology, 432(9), 2930–2948. https://doi.org/10.1016/j.jmb.2019.12.027

García de la Torre, J., Huertas, M. L., & Carrasco, B. (2000). HYDRONMR: Prediction of NMR Relaxation of Globular Proteins from Atomic-Level Structures and Hydrodynamic Calculations. Journal of Magnetic Resonance, 147(1), 138–146. https://doi.org/10.1006/jmre.2000.2170

Halle, B. (2004). Protein hydration dynamics in solution: a critical survey. Philosophical Transactions of the Royal Society of London. Series B: Biological Sciences, 359(1448), 1207–1224. https://doi.org/10.1098/rstb.2004.1499

Halle, B., & Davidovic, M. (2003). Biomolecular hydration: From water dynamics to hydrodynamics. Proceedings of the National Academy of Sciences, 100(21), 12135–12140. https://doi.org/10.1073/pnas.2033320100

Harding, S. E. (1997). The intrinsic viscosity of biological macromolecules. Progress in measurement, interpretation and application to structure in dilute solution. Progress in Biophysics and Molecular Biology, 68(2-3), 207–262. https://doi.org/10.1016/S0079-6107(97)00027-8

Harding, S. E. (2001). The hydration problem in solution biophysics: an introduction. In Biophysical Chemistry (Vol. 93).

Hellman, L. M., Rodgers, D. W., & Fried, M. G. (2010). Phenomenological partial-specific volumes for G-quadruplex DNAs. European Biophysics Journal, 39(3), 389–396. https://doi.org/10.1007/s00249-009-0411-7

Hwang, C.-C., Chang, P.-R., Hsieh, C.-L., Chou, Y.-H., & Wang, T.-P. (2019). Thermodynamic analysis of remote substrate binding energy in 3α-hydroxysteroid dehydrogenase/carbonyl reductase catalysis. Chemico-Biological Interactions, 302, 183–189. https://doi.org/10.1016/j.cbi.2019.02.011

Kauzmann, W. (1959). Some Factors in the Interpretation of Protein Denaturation. In Advances in Protein Chemistry (Vol. 14, pp. 1–63). https://doi.org/10.1016/S0065-3233(08)60608-7

Kovacic, R. T., & van Holde, K. E. (1977). Sedimentation of homogeneous double-strand DNA molecules. Biochemistry, 16(7), 1490–1498. https://doi.org/10.1021/bi00626a038

Kuntz, I. D. (1971). Hydration of Macromolecules. III. Journal of the American Chemical Society, 93(2), 514–516.

Kuntz, I. D., & Kauzmann, W. (1974). Hydration of Proteins and Polypeptides. Advances in Protein Chemistry, 28, 239–345. https://doi.org/10.1016/S0065-3233(08)60232-6

Lee, J. C., & Timasheff, S. N. (1977). In vitro reconstitution of calf brain microtubules: effects of solution variables. Biochemistry, 16(8), 1754–1764. https://doi.org/10.1021/bi00627a037

Li, J. (2005). Not so crystal clear: the structure of the human telomere G-quadruplex in solution differs from that present in a crystal. Nucleic Acids Research, 33(14), 4649–4659. https://doi.org/10.1093/nar/gki782

Perrin, F. (1936). Mouvement Brownien d’un ellipsoide (II). Rotation libre et dépolarisation des fluorescences. Translation et diffusion de molécules ellipsoidales. Journal de Physique et Le Radium, 7(1), 1–11. https://doi.org/10.1051/jphysrad:01936007010100

Persson, F., Söderhjelm, P., & Halle, B. (2018). How proteins modify water dynamics. Journal of Chemical Physics, 148(21). https://doi.org/10.1063/1.5026861

Rocco, M., & Byron, O. (2015). Hydrodynamic Modeling and Its Application in AUC. Methods in Enzymology, 562, 81–108. https://doi.org/10.1016/bs.mie.2015.04.010

Rowe, A. J. (1977). The concentration dependence of transport processes: A general description applicable to the sedimentation, translational diffusion, and viscosity coefficients of macromolecular solutes. Biopolymers, 16(12), 2595–2611. https://doi.org/10.1002/bip.1977.360161202

Rupley, J. A., & Careri, G. (1991). Protein Hydration and Function. Advances in Protein Chemstry, 41, 37–172. https://doi.org/10.1016/S0065-3233(08)60197-7

Rupley, J. A., Gratton, E., & Careri, G. (1983). Water and globular proteins. Trends in Biochemical Sciences, 8(1), 18–22. https://doi.org/10.1016/0968-0004(83)90063-4

Sengupta, N., Jaud, S., & Tobias, D. J. (2008). Hydration Dynamics in a Partially Denatured Ensemble of the Globular Protein Human α-Lactalbumin Investigated with Molecular Dynamics Simulations. Biophysical Journal, 95(11), 5257–5267. https://doi.org/10.1529/biophysj.108.136531

Squire, P. G., & Himmel, M. E. (1979). Hydrodynamics and protein hydration. Archives of Biochemistry and Biophysics, 196(1), 165–177. https://doi.org/10.1016/0003-9861(79)90563-0

Stafford, W. F. (2016). Analysis of Nonideal, Interacting, and Noninteracting Systems by Sedimentation Velocity Analytical Ultracentrifugation. In Analytical Ultracentrifugation (pp. 463–482). Springer Japan. https://doi.org/10.1007/978-4-431-55985-6_23

Stafford, W. F., & Sherwood, P. J. (2004). Analysis of heterologous interacting systems by sedimentation velocity: curve fitting algorithms for estimation of sedimentation coefficients, equilibrium and kinetic constants. Biophysical Chemistry, 108(1-3), 231–243. https://doi.org/10.1016/j.bpc.2003.10.028

Tarek, M., & Tobias, D. J. (2008). The role of protein–solvent hydrogen bond dynamics in the structural relaxation of a protein in glycerol versus water. European Biophysics Journal, 37(5), 701–709. https://doi.org/10.1007/s00249-008-0324-x

Umehara, T., Kuwabara, S., Mashimo, S., & Yagihara, S. (1990). Dielectric study on hydration of B-, A-, and Z-DNA. Biopolymers, 30(7-8), 649–656. https://doi.org/10.1002/bip.360300702

Vulevic, B., & Correia, J. J. (1997). Thermodynamic and structural analysis of microtubule assembly: the role of GTP hydrolysis. Biophysical Journal, 72(3), 1357–1375. https://doi.org/10.1016/S0006-3495(97)78782-4

Williams, M. A., Goodfellow, J. M., & Thornton, J. M. (1994). Buried waters and internal cavities in monomeric proteins. Protein Science, 3(8), 1224–1235. https://doi.org/10.1002/pro.5560030808

Wright, R. T., Hayes, D. B., Stafford, W. F., Sherwood, P. J., & Correia, J. J. (2018). Characterization of therapeutic antibodies in the presence of human serum proteins by AU-FDS analytical ultracentrifugation. Analytical Biochemistry, 550, 72–83. https://doi.org/10.1016/j.ab.2018.04.002

Wüthrich, K., Billeter, M., Güntert, P., Luginbühl, P., Riek, R., & Wider, G. (1996). NMR studies of the hydration of biological macromolecules. Faraday Discuss., 103, 245–253. https://doi.org/10.1039/FD9960300245

Wuttke, R., Hofmann, H., Nettels, D., Borgia, M. B., Mittal, J., Best, R. B., & Schuler, B. (2014). Temperature-dependent solvation modulates the dimensions of disordered proteins. Proceedings of the National Academy of Sciences, 111(14), 5213–5218. https://doi.org/10.1073/pnas.1313006111

Yang, D., Correia, J. J., Stafford III, W. F., Roberts, C. J., Singh, S., Hayes, D., Kroe-Barrett, R., Nixon, A., & Laue, T. M. (2018). Weak IgG self-and hetero-association characterized by fluorescence analytical ultracentrifugation. Protein Science, 27(7), 1334–1348. https://doi.org/10.1002/pro.3422

Yang, P. H., & Rupley, J. A. (1979). Protein-water interactions. Heat capacity of the lysozyme-water system. Biochemistry, 18(12), 2654–2661.

Zhou, H.-X. (1995). Calculation of Translational Friction and Intrinsic Viscosity. II. Application to Globular Proteins. Biophysical Journal, 69.

