## Supplemental Information for "Revisiting Macromolecular Hydration with HullRadSAS"

**SUPPLEMENTAL FIGURE S1** HullRadSAS and HullRad predict the same translational hydrodynamic radii.

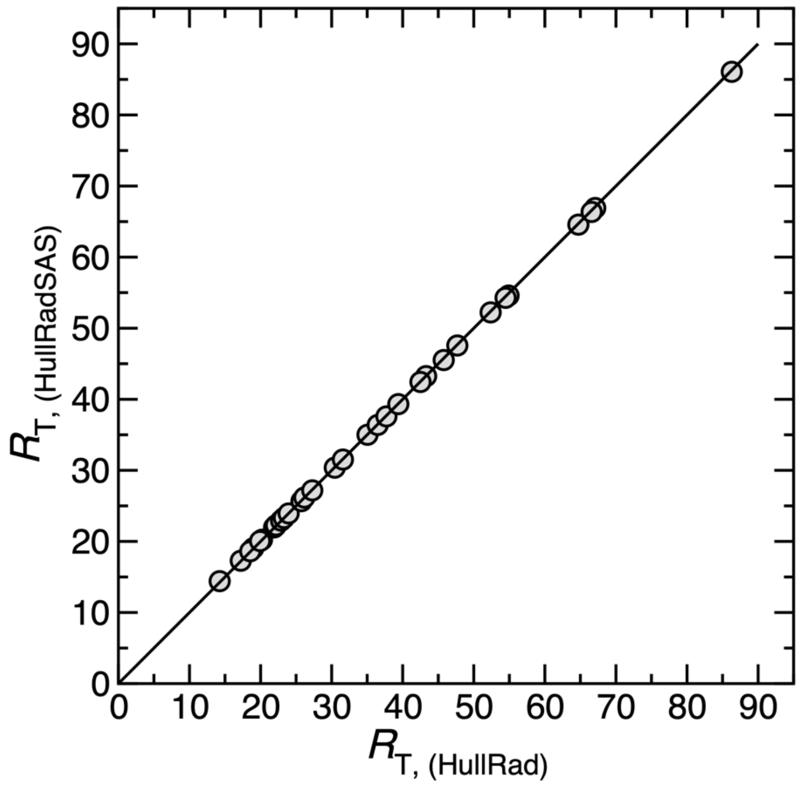

HullRadSAS predicted $R_T$ values for folded proteins are plotted versus the HullRad predicted values for the data set of proteins listed in Supplemental Table S1. The black line through the data points represents a slope of one and intercept of zero.

**SUPPLEMENTAL FIGURE S2** Optimization of the SAS probe radius.

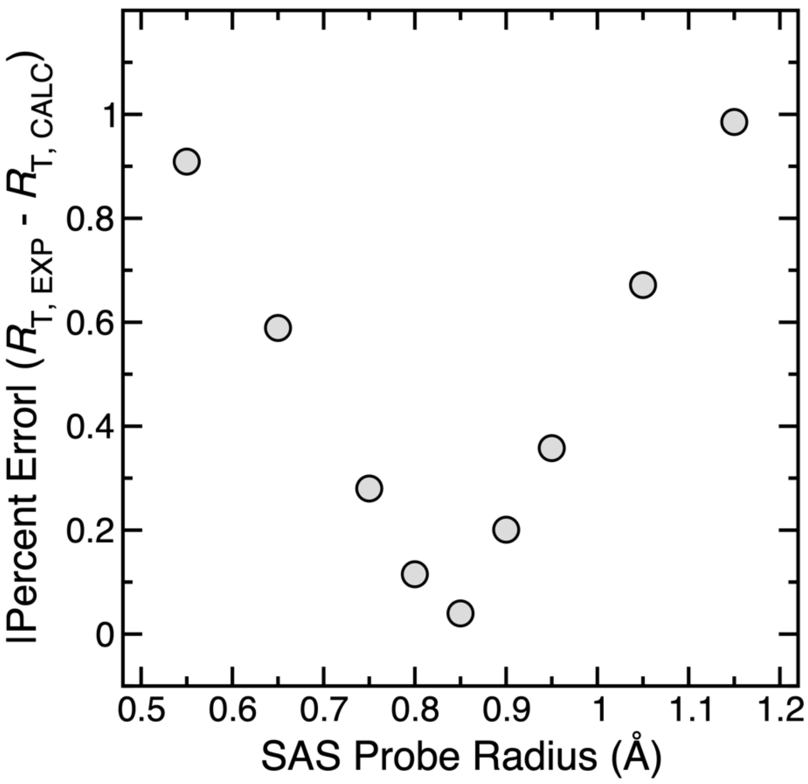

Translational hydrodynamic radii ($R_T$) were calculated using HullRadSAS for the proteins listed in Supplemental Table S1. The absolute value percent error between the experimental value and calculated value of $R_T$ for different SAS probe radii is plotted as gray circles. An optimal probe radius of 0.85 Å was chosen.

**SUPPLEMENTAL FIGURE S3** HullRadSAS accurately predicts intrinsic viscosities.

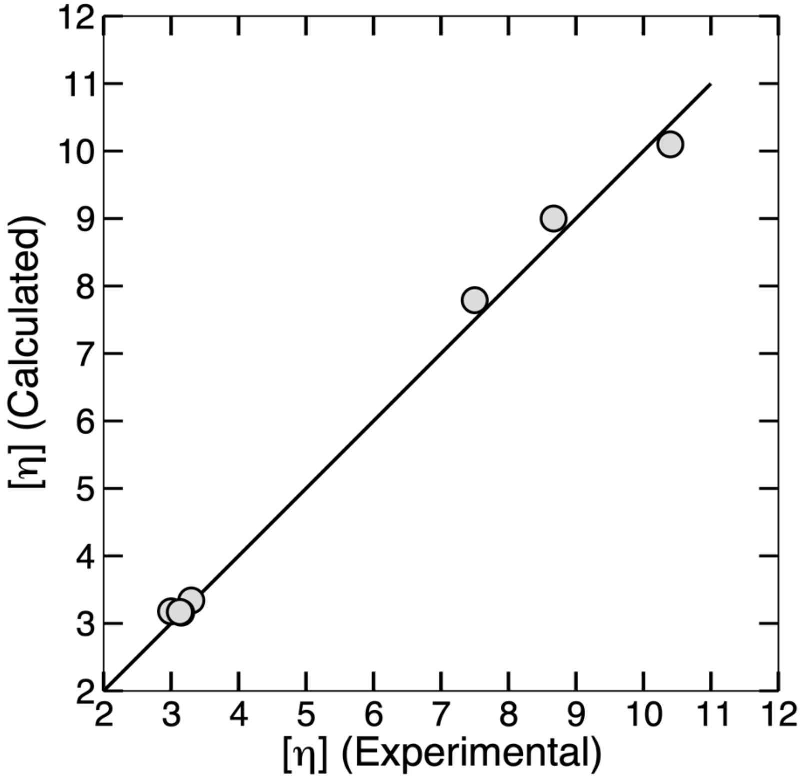

Intrinsic viscosities [η] calculated using HullRadSAS are plotted versus experimental values for the proteins and DNA listed in Supplemental Table S2. The black line through the data points represents a slope of one and intercept of zero.

**SUPPLEMENTAL TABLE S1** Translational Hydrodynamic Radii for Folded Proteins.

| Protein (PDB Code) | MW (Da) | $R_{T,Exp}$[a] (Å) | $R_{T,HR}$[b] (Å) | $R_{T,HRSAS}$[c] (Å) |
|---|---|---|---|---|
| BPTI (5PTI) | 6518 | 14.5 | 14.3 | 14.4 |
| Cytochrome *c* (1HRC) | 11703 | 17.3 | 17.2 | 17.3 |
| RNase A (8RAT) | 13692 | 18.9 | 19.0 | 19.0 |
| Lysozyme (1AKI) | 14315 | 18.6 | 18.5 | 18.6 |
| LegHb (1LH1) | 16657 | 21.4 | 20.2 | 20.2 |
| Apo-Mb (1MGN) | 17359 | 20.8 | 19.9 | 20.0 |
| Soy TI (1AVU) | 19984 | 22.6 | 21.9 | 22.0 |
| Trypsinogen (1TGN) | 22620 | 22.1 | 22.1 | 22.2 |
| β-Trypsin (1TPO) | 23124 | 22.8 | 22.2 | 22.3 |
| α-Chymotrypsin (4CHA) | 25041 | 22.7 | 22.9 | 22.9 |
| Chymotrypsinogen (2CGA) | 25670 | 22.6 | 23.3 | 23.3 |
| Carbonic Anhydrase (2CAB) | 28757 | 24.1 | 23.9 | 23.9 |
| Superoxide Dismutase (2SOD) | 31089 | 25.9 | 25.8 | 25.7 |
| Pepsin (4PEP) | 34516 | 24.6 | 26.2 | 26.2 |
| β-Lactoglobulin (1BEB) | 36606 | 27.4 | 27.3 | 27.2 |
| TPI (8TIM) | 52996 | 29.7 | 30.5 | 30.4 |
| Hb (CO) (1HCO) | 61942 | 31.5 | 31.6 | 31.5 |
| HSA (1AO6) | 66482 | 34.0 | 35.1 | 35.0 |
| Alkaline Phosphatase (1ALK) | 94082 | 37.6 | 37.0 | 36.4 |
| Citrate Synthase (1CTS) | 97835 | 37.0 | 37.7 | 37.6 |
| Inorganic PPase (2AU9) | 117361 | 37.6 | 39.4 | 39.3 |
| Tryp. Synthase (1KFK) | 138595 | 44.4 | 45.8 | 45.5 |
| Human IgG (1HZH) | 143337 | 55.1 | 55.0 | 54.6 |
| Apo G3PD (2CG1) | 143743 | 42.9 | 43.3 | 43.3 |
| Apo LDH (5LDH) | 145749 | 42.5 | 42.5 | 42.4 |
| Aldolase (1ADO) | 156776 | 47.6 | 47.7 | 47.6 |
| Holo Catalase (4BLC) | 230321 | 52.3 | 52.4 | 52.2 |
| Xanthine Oxidase (1FIQ) | 267770 | 54.5 | 54.5 | 54.3 |
| β-Galactosidase (4V40) | 464490 | 68.5 | 67.1 | 66.9 |
| Apo-Ferritin (3AJO) | 512084 | 67.4 | 64.7 | 64.6 |
| Urease (3LA4) | 539700 | 65.8 | 66.6 | 66.4 |
| GroEL (2CGT) | 828989 | 82.8 | 86.3 | 86.1 |

[a]Experimental $R_T$ values from Table 1 of Fleming and Fleming (Fleming & Fleming, 2018).
[b]$R_T$ calculated with HullRad.
[c]$R_T$ calculated with HullRadSAS.

**SUPPLEMENTAL TABLE S2** Intrinsic Viscosities for Folded Proteins and DNA Duplexes.

| Protein (PDB Code) | MW (kDa) | $[\eta]_{exp}$ (ml/g) | $[\eta]_{calc}^{a}$ (ml/g) | $[\eta]_{calc}$-$[\eta]_{exp}$ (%Error) | $[\eta]_{exp}$ Reference |
|---|---|---|---|---|---|
| RNase A (8RAT) | 13692 | 3.30 | 3.34 | 1.3 | (Buzzell & Tanford, 1956) |
| Lysozyme (1AKI) | 14315 | 3.00 | 3.18 | 6.0 | (Sophianopoulos et al., 1962) |
| Apo-Mb (1MGN) | 17359 | 3.15 | 3.16 | 0.3 | (Wyman & Ingalls, 1943) |
| Chymotrypsinogen (2CGA) | 25670 | 3.13 | 3.17 | 1.4 | (Schwert, 1951) |
| Human IgG (1HZH) | 143337 | 8.67 | 9.00 | 3.8 | (Monkos & Turczynski, 1999) |
| B-DNA 20mer | 12270 | 7.50 | 7.79 | 3.9 | (Tsortos et al., 2011) |
| B-DNA 30mer | 19129 | 10.40 | 10.10 | -2.9 | (Tsortos et al., 2011) |

[a]Calculated values from HullRadSAS

**REFERENCENCES**

Buzzell, J. G., & Tanford, C. (1956). The Effect of Charge and Ionic Strength on the Viscosity of Ribonuclease. *The Journal of Physical Chemistry*, *60*(9), 1204–1207. https://doi.org/10.1021/j150543a014

Fleming, P. J., & Fleming, K. G. (2018). HullRad: Fast Calculations of Folded and Disordered Protein and Nucleic Acid Hydrodynamic Properties. *Biophysical Journal*, *114*(4), 856–869. https://doi.org/10.1016/j.bpj.2018.01.002

Monkos, K., & Turczynski, B. (1999). A comparative study on viscosity of human, bovine and pig IgG immunoglobulins in aqueous solutions. *International Journal of Biological Macromolecules*, *26*(2–3), 155–159. https://doi.org/10.1016/S0141-8130(99)00080-X

Schwert, G. W. (1951). The Molecular Size and Shape of the Pancreatic Proteases. *Journal of Biological Chemistry*, *190*(2), 799–806. https://doi.org/10.1016/S0021-9258(18)56030-0

Sophianopoulos, A. J., Rhodes, C. K., Holcomb, D. N., & van Holde, K. E. (1962). Physical Studies of Lysozyme. *Journal of Biological Chemistry*, *237*(4), 1107–1112. https://doi.org/10.1016/S0021-9258(18)60292-3

Tsortos, A., Papadakis, G., & Gizeli, E. (2011). The intrinsic viscosity of linear DNA. *Biopolymers*, *95*(12), 824–832. https://doi.org/10.1002/bip.21684

Wyman, J., & Ingalls, E. N. (1943). A Nomographic Representation of Certain Properties of the Proteins. *Journal of Biological Chemistry*, *147*(2), 297–318. https://doi.org/10.1016/S0021-9258(18)72384-3
